# Brain volumes of very low birth weight infants measured by two dimensional cranial ultrasonography, a prospective cohort study

**DOI:** 10.1101/485474

**Authors:** Gülsüm Kadıoğlu Şimşek, Fuat Emre Canpolat, Mehmet Büyüktiryaki, Gözde Kanmaz Kutman, Cüneyt Tayman

**Affiliations:** Department of Neonatology, NICU, Zekai Tahir Burak Health Practice and Research Center, University of Medical Sciences 06240, Hamamönü /Altındağ Ankara TURKEY

**Keywords:** cranial ultrasonography, brain volume, ventricular volume, very low birth weight, preterm newborn infant

## Abstract

**Background:** Cranial ultrasonography is the main neuroimaging technique for very low birth weight infants. Brain volume is a very important information about central nervous system of preterm babies.

This study aimed to evaluate brain volumes of preterm infants with two dimensional measurements of cranial ultrasonography.

**Methods:** Intracranial height, anteroposterior diameter, bi-parietal diamater, ventricular height, thalamo-occipital distance and ventricular index measured with routine cranial ultrasonographic scanning. Brain considered a spheric, ellipsoid model and estimated absolute brain volume (EABV) calculated by substracting two lateral ventricular volumes from the total brain volume.

**Results:** One hundred and twenty one preterm infants under a birthweight of 1500 g and 32 weeks of gestational age included in this study. Mean gestational age of study population was 27,7 weeks, and mean birthweight was 1057 grams.

Twenty two of 121 infants had dilated ventricle, in this group EABV was lower than normal group (202 ± 58 cm^3^ vs 250 ± 53 cm^3^, respectively, p<0.01). Advanced resuscitation, bronchopulmonary dysplasia and late onset sepsis found to be independent risk factors for low brain volume in our data.

**Conclusions:** Estimated absolute brain volume could be calculated and estimated by two dimensional measurements with transfontanel ultrasonography.

## Background

Nonhemorrhagic ventriculomegaly (NHVM) or dilated ventricle is a common finding in very preterm infants (1). Nonhemorrhagic ventriculomegaly is associated with increased risks of neurodevelopmental problems among these preterm newborns (2). Although exact measurements have been used, there is no concensus on which measurements or cutoff values should be applied. This problem is about preterm infants who do not experience cranial bleeding or periventricular leukomalacia (PVL) because this pathologies should be evaluated seperately. Previous studies showed that the ventricular-brain ratio is another way of estimation brain volume in association with head circumference and widths of lateral ventricular horns (1). There is some evidence about a good correlation between ultrasonographic measurements and magnetic resonance imaging for volume of ventricles for preterm infants (3). Brain volume of preterm infants is an important factor that has an association with neurodevelopmental outcome (4). Magnetic resonance imaging (MRI) or computerized tomography is not a routine advanced imaging technique for preterm infants without intraventricular hemorrhage (IVH), PVL and/or additional brain lesions (5,6). Furthermore transfontanel ultrasonography is the simplest method to assess preterm brain, appropriate for scanning, easy to use, accessible and even could be found in low resource settings (5). We hypothesized that we can calculate the absolute brain volume by subtracting the ventricular volumes from the total brain volume measured and calculated with two dimensional (2D) distances by ultrasonography without advanced neuroimaging. Consequently the main aim of this study was to measure 2D distances obtained by ultrasonography and estimate absolute brain volume.

## Methods

This prospective study was conducted in Neonatal Intensive Care Unit (NICU) at University of Health Sciences Zekai Tahir Burak Health, Practice and Research Center Hospital, Ankara Turkey. This perinatal center has a NICU with 130 beds and 18,000 birth per year. During the study period (January 2017-May 2018) 341 infants were born under 1500 gr and 32 weeks of gestational age. Of these 341 infants, 55 died within the first week, 41 were small for gestational age, 72 had Grade III-IV cranial bleeding, 12 had prominent periventricular leukomalacia (PVL), 40 patient dropped out because of, lost follow up (18), declined participation (14) and died (8) before the last examination (34 weeks of gestational age). A total of remaining 121 infants were eligible for all serial cranial ultrasound examinations (Figure 1). Local review board ratification (Medical Specialty Training Board, TUEK) and Ethical Committe Approval obtained for this study.

**Figure 1.**
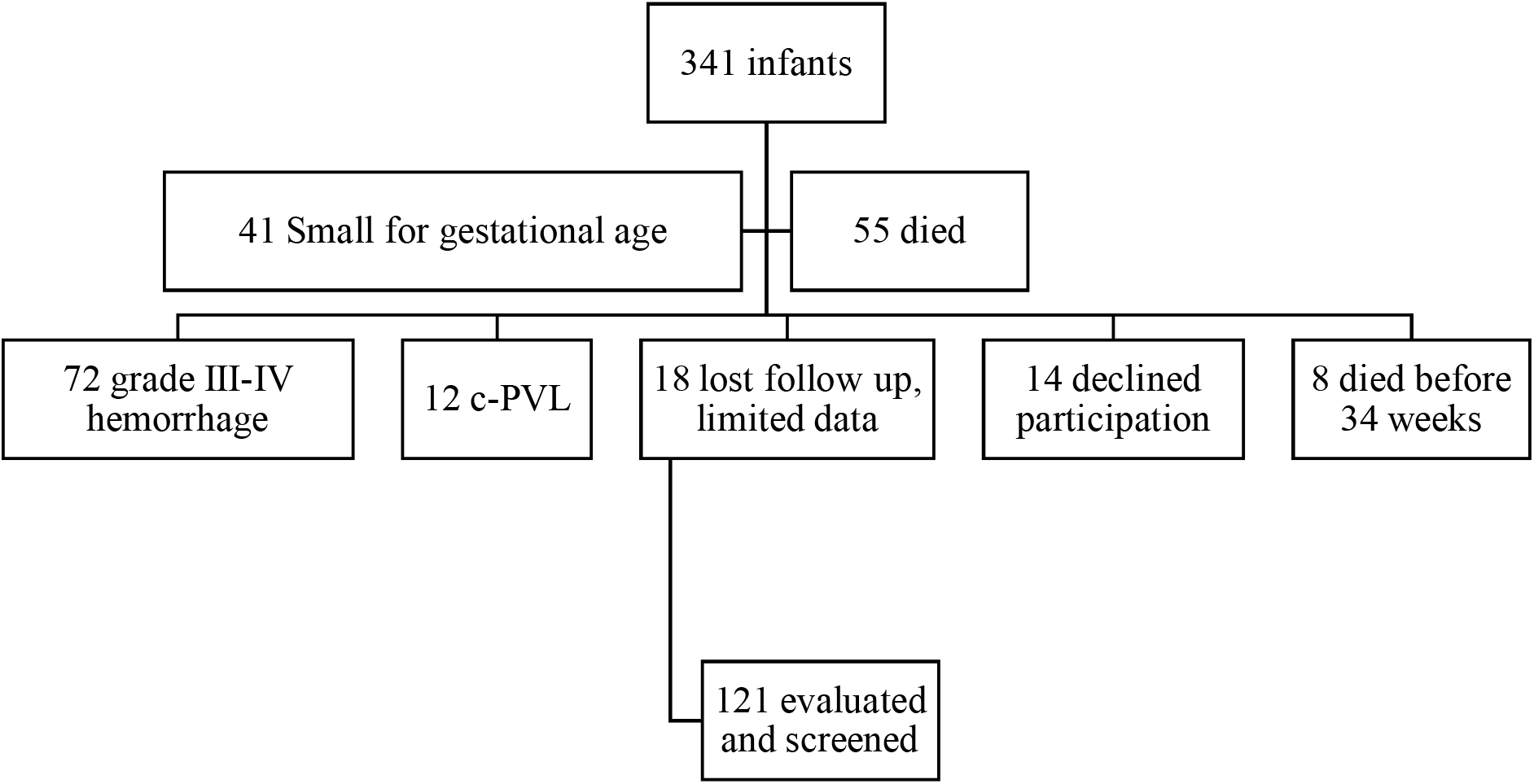
Flow chart of study population

### Clinical Definitions

Advanced resuscitation in delivery room, if an infants need intubation, and/or chest compression, or drug administration in the delivery room soon after birth.

Prolonged parenteral nutrition means if an infant needs parenteral nutrition more than 2 weeks.

Delayed regain of birthweight, if an infant reaches birthweight for more than 2 weeks.

Feeding intolerance is consist of delayed meconium passage, significant abdominal distension without significant emesis or orogastric tube output but more than >50% gastric residuals at least two times a day, as a result feeding intolerance cause a feeding break.

Late onset sepsis (culture proven) defined as, clinical worsening with skin abnormalities, gastrointestinal dysfunctions, bradycardia or tachycardia, hypo-hyperthermia, apnea, and hematological (leukopenia, immature-mature neutrophyl ratio more than 0.2 in differentials) and biochemical (elevated C-reactive protein and/or interleukin-6, hypo-hyperglycemia) findings of sepsis after 7 days of postnatal age without a blood culture positivity (6).

Bronchopulmonary dysplasia, defined as moderate and severe forms of the disease described before as oxygen dependence at 36 weeks of postmenstrual age (7).

Other morbidities such as respiratory distress syndrome, surfactant administration, patent arterial duct, hypotension, days passed on ventilator, necrotizing enterocolitis, anemia, blood transfusions, postnatal steroids, retinopathy of prematurity etc. also noted and entered to the database.

### Cranial ultrasonographic examinations

Cranial ultrasound was performed (Toshiba^®^ Aplio^™^ 300, Canon^®^ Medical Systems Turkey, with a 7 Mhz vector transducer probe) by the attending skilled neonatologist for all infants once within postnatal 3-5 days, on the second week, thereafter monthly and at least once at 34 weeks of corrected age. Routine radiologic cranial ultrasonographic examinations were also made by consultant radiologist unaware of this study. Radiologists did not measured all measurements as we did except ventricular index. We have to do our last scanning in NICU at 34 weeks of gestational age because most of the patients were discharged near 34th week. Ultrasound was focused on finding IVH (grading according to Papile et al,[8]) and PVL. Cystic periventricular leukomalacia was defined as multiple cysts with a typical region in the posterior periventricular white matter near to lateral aspect of the trigone of lateral ventricle and in the white matter adjacent to the foramen of Monro. Definition of non-hemorrhagic ventriculomegaly is a ventricular index (9) should not be larger than 97 percentile+ 4mm and larger than 90 p. Previous studies showed that a linear measurement of VI for a preterm infant at 34 corrected age 97p is 12 mm (10,11).

### Calculation of brain volume

The three cerebral caliber considered for the model (Figure 2A) were half of the intracranial bi-parietal diameter (R1 = BPDia / 2), intracranial postero-anterior or fronto-occipital diameter (R2 = APD / 2) and cranial height (R3 = ICH / 2). Bone and extracerebral space were not included in these measurements in our study. The transvers plane of the cranium at the level of the maximal occipital-frontal circumference was considered an ellipse for area estimation (area = π X R1 X R2); the cranium was considered as an ellipsoid for volume estimation (volume = 4/3 X π X R1 X R2 X R3)(12).

**Figure 2.**
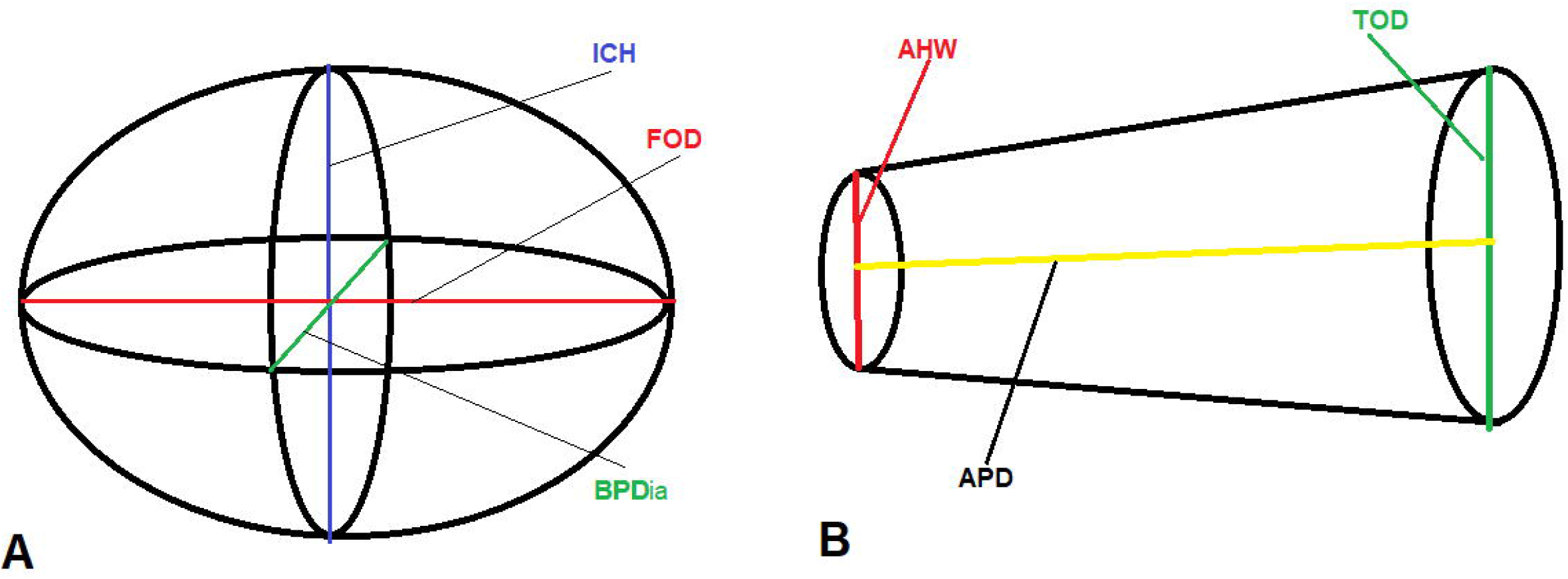
Measurements of brain and ventricles as geometric figures. ICH, Intracranial height; BPDia, Bi-parietal diamater; APD, FOD, Anteroposterior diamater or fronto-occipital distance; AHW, Anterior horn width; TOD, Thalamo-occipital distance

### Calculation of ventricular volume

Lateral ventricles are reverse “C” shaped, asymmetric formations. Ventricular volume can be measured by 2D or 3D dimensional ultrasound (3,13,14) and magnetic resonance imaging. We calculated estimated ventricular volume as calculating volume of a truncated cone which is confirmed mathematically. The small radii is represented by anterior horn width (AHW), the big radii was thalamo-occipital distance (TOD) and the height of the cone was APD (Figure 2B). Each ventricle measured and calculated seperately beacuse of assymetry. All measured distances are explained in Table 1 and Figure 3. Estimated absolute brain volume (EABV) obtained by substracting two lateral ventricular volumes from the brain volume.

**Table 1.** Description of linear cranial ultrasound measurements.

| Measurement | Plane | Description | Reference |
| --- | --- | --- | --- |
| Bi-parietal Diameter (BPDia) | Coronal (foramen of Monro) | Intracranial bi-parietal diameter measured at the level of the Sylvian fissures | 10 |
| Antero-posterior Diameter (APD) | Mid-sagittal | Intracranial occipito-frontal diameter(similar reference points to the ones used for head circumference measurement) | 10 |
| Cranial height (ICH) | Mid-sagittal | Intracranial height from the posterior aspect of the foramen magnum to the inner aspect of the fontanel below the transducer | 10 |
| Thalamo-occipital Distance (TOD) | Parasagittal (atrium) | Measured from the most posterior point of the thalamus to the furthest depth of the occipital horn posteriorly | 10,11 |
| Anterior horn width (AHW) | Coronal (foramen of Monro) | Maximum distance within the ventricle | 10,11 |
| Levene ventricular Index (VI) | Coronal (foramen of Monro) | Horizontal distance between the external tip of the lateral ventricle and the midline | 11,12 |

|  | Normal<br>n=99 | Dilated ventricle<br>n=22 | p |
| --- | --- | --- | --- |
| Gestational age, weeks | 27 ± 2 | 28 ± 2,8 | 0,269 |
| Birthweight, grams | 1058 ± 226 | 1052 ± 269 | 0,910 |
| CRIB-II score, median | 5 | 4 | 0.558* |
| EABV, cm<sup>3</sup> | 250 ± 53 | 202 ± 58 | 0.001 |
| VI, mm | 9,7 ± 1.9 | 12,8 ± 1,7 | 0,0001 |
| VH, mm | 3 ± 1 | 5 ± 2 | 0,001 |
| BPDia,mm | 71 ± 7 | 76 ± 7 | 0,02 |

**Figure 3.**
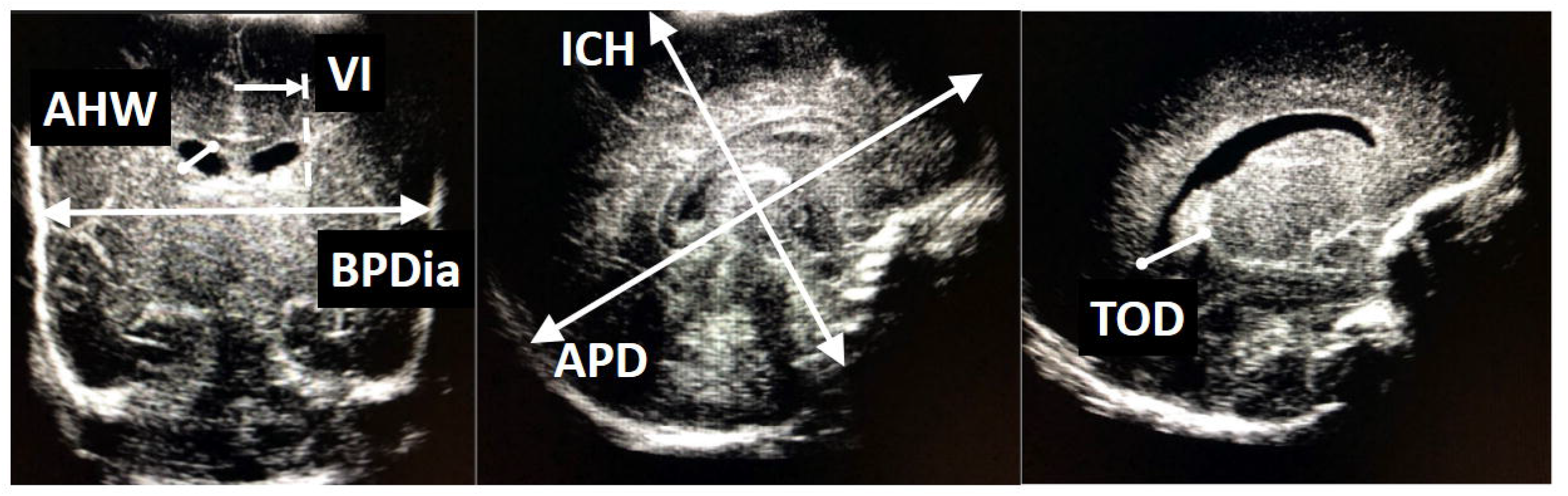
Measurements of brain and ventricles on ultrasonographic images. ICH, Intracranial height; BPDia, Bi-parietal diamater; APD, FOD, Anteroposterior diamater or fronto-occipital distance; AHW, Anterior horn width; TOD, Thalamo-occipital distance; VI,ventricular index

### Statistical analysis

All this data entered a personal computer, by using MS Excel® and with its ‘function’ property. Then all these data entered to SPSS for Windows® version 21 to assess statistical analysis. Ranges, percentiles, descriptives and frequencies used for distributions, Multi nominal logistic regression used for dilated ventricle as dependent variable, covariates designed as birthweight and gestational age, independent variables were, resuscitation in the delivery room, prolonged parenteral nutrition, delay of regain birthweight, feeding intolerance, late onset sepsis and bronchopulmonary dysplasia which were all statistically different between groups (infants with and without NHVM) shown by Chi-square test. Pearson correlation test also used for cranial measurements.

## Results

Mean gestational age of study population (n=121) was 27,7 weeks (23-31), and mean birthweight was 1057 g (500-1490). Fifty four (44,6%) of 121 were male, 94 (77%) were delivered via cesarean section. Mean CRIB score was 4 (1-13), 22 (18,2 %) had dilated ventricle, 51 (42%) of these infants had advanced resuscitation, 50 (41%) received parenteral nutrition more than two weeks, 49 (40%) infants reached their birthweights more than two weeks, 48 (39%) had some experience about feeding intolerance, 54 (44%) had clinically suspected late onset sepsis, 40 (33%) had moderate to severe bronchopulmonary dysplasia. We also compared clinical characteristics of infants with and without dilated ventricle (Table 2). After adjusting data with gestational age and birthweight, resuscitation, bronchopulmonary dysplasia and late onset sepsis were independent risk factors for low brain volume and NHVD (Table 3). Respiratory distress syndrome, surfactant administration, patent arterial duct, hypotension, days on mechanical ventilator, necrotizing enterocolitis, anemia, blood transfusions, postnatal steroids, retinopathy of prematurity do not independently associate to ventricular enlargement and low brain volume in our study population.

**Table 2.** Comparison of infants with and without dilated ventricle.

|  |  |  |  |
| --- | --- | --- | --- |
| ICH, mm | 65±10 | 73± 6 | 0,001 |
| TOD, mm | 9 ±2 | 12±4 | 0.001 |

\*non-parametric test,
CRIB, Clinical risk index for babies; EABV, estimated absolute brain volume; VI, ventricular index; VH, ventricular height, BPDia, Biparietal diameter; ICH, intracranial height; TOD, Thalamo-occipital distance

**Table 3.** Risk factors affecting non-haemorrhagic ventricular dilatation.

| **Risk factors** | **OR** | **95 % CI** | **p** |
| --- | --- | --- | --- |
| **Resuscitation** | **2,9** | **1,1-7,65** | **0,028\*** |
| Prolonged parenteral nutrition | 2,4 | 0,94-6,21 | 0,06 |
| Delayed weight gain | 2,0 | 0,79-5,11 | 0,142 |
| Feeding intolerance | 1,6 | 0,66-4,24 | 0,276 |
| **Bronchopulmonary dysplasia** | **4,2** | **1,5-11,8** | **0,006\*** |
| **Late onset sepsis** | **5,3** | **1,8-15,8** | **0,002\*** |

\*Statistically significant

Biparietal diameter and ICH were highly correlated with brain volume (r: 0.854, p:0.001, r:0.845, p:0.001 respectively).

## Discussion

To the best of our knowledge this is the first study which calculates and estimates absolute brain volume with 2D cranial ultrasound measurements in very low birth weight infants. Our data suggests that brain volume could be calculated by 2D ultrasonographic distances and this is the most easiest way to observe preterm infant’s estimated absolute brain volume. Even in low resource settings, cranial ultrasonography is the main neuroimaging method that has low cost, easy to use, safe and simple. Ventriculomegaly alone is an important issue for preterm infants and associated with poor neurologic outcome (2). We can say that “the larger ventricle, the smaller brain tissue” and this is associated with poorer neurologic outcome (15) so that brain tissue volume of preterm infants is also another way of predicting neurologic outcome.

In many studies, low brain volume is associated with morbidites and neurodevelopmental impairment (16,17). All preterm infants should be evaluated for cranial pathologies, especially ventricular hemorrhage, PVL and dilated ventricles (NHVM). Ultrasonographic screening is recommended for preterm infants under 30 weeks of gestational once between 7 and 14 days of age and should be optimally repeated between 36 and 40 weeks’ postmenstrual age and this is recommended by American Academy of Pediatrics, American Academy of Neurology and many other national medical societies (5). There is no recommendation that all preterms under 1500 gr or less than 32 weeks of gestational age should undergo advanced neuroimaging such as cranial tomography and/or cranial MRI except additional clues like pathologic neurologic examination, ventricular bleeding, hydrocephalus, congenital malformations of central nervous system and PVL. Therfore very preterm infants with less morbidites and minor complications do not undergo routine advanced neuroimaging in many NICUs but they usually have at least a cranial US scan.

The ventricular index is one of the important parameter to diagnose or scan ventricular dilation. Previous studies published nomogram for the VI in newborns (9,18). Both data showed that upper limit of VI depends on gestational week. The reference values of these studies correlate well for term and near term neonates, while those for very low birth weight newborns show more variation. This may be associated that the lower the post-menstrual age the fewer infants were studied, increase the error of mean. Liao et al. studied preterm infants near 26-27 weeks of gestation, who are at risk of developing ventricular dilation (18). Additionally only evaluation of anterior horn and VI is not enough to say ventricular dilation or reduced brain volume. Therefore more data and new techniques are needed especially for VLBW infants.

Another important measurement is anterior horn width (AHW). In the majority of newborns, the AHW is less than 3 mm (11,18-20). The clinical importance of an AHW more than 3 mm in the absence of bleeding is not obvious. An AHW between 3 and 5 mm was not associated with neurodevelopmental impairment at a follow up visit in a group of thirteen babies within the first year of life (20). Values over 6 mm, however, are associated with ventricular distension and suggest the need for surgical treatment according to Govaert et al. and may be associated with reduced brain volume in the absence of cranial-ventricular hemorrhage (21). The other main measurement of preterm ventricle is TOD. Controversy also exists whether normal values of occipital horn size and TOD depend on gestational week. Furthermore, the variation in reported reference curves is remarkable (18). Reeder and friends reported that (22) regardless of the degree of prematurity, an occipital horn length larger than 16 mm suggests main cerebral problmes in premature infants. Davies and friends (11) reported much higher reference ranges for TOD in preterm neonates, with an upper limit of 24.7 mm.

One of the other issue about brain measurements of preterm infant is ventricular asymmetry. During brain growth ventricular asymmetry may be infulenced by genetic and environmental factors (10,23). It has also been documented that in case of ventricular asymmetry, ventricle dilation is frequently associated with a bigger ipsilateral choroid plexus. In previous studies, relationship was found between asymmetric ventricle and mode of delivery, gender and the infants age at the time of cranial ultrasonography (10,23).

Extracerebral space is another parameter should be taken into consideration for brain volume. Some infants have large extracerebral space in our study but we did not calculate additional volume for extracerebral space. In our study, intracranial measurements started at the margin of cortical brain tissue as shown in the figures.

Ventricular/brain ratio and lateral ventricular diameter are also investigated for assessing brain volume and predicting neurologic outcome (1). The authors concluded that the ventricular/ brain ratio, size of the lateral ventricles, and head circumference are suitable measures for the estimation of total and regional brain volumes and this study confirmed that ventriculomegaly is strongly associated with cerebral pathologies (1).

As we observed that each of measurements are associated with ventricular dilation seperately and all have some predictive power for assessing brain volume. We hypothesized that a calculation which covers all these parameters will give a more proper information about preterm brain volume. We suggest that sonographers and clinicans should make all these measurements as they do before and enter these data to a computer based system and could calculate and estimate brain volumes of very preterm infants easily.

Of course there are some limitations of this study. One of them is we could not confirm these calculations by MRI or similar advanced neuroimaging techniques. But this should not be a major shortcoming because further studies and measurements may be established with these ultrasonographic measurements and next years more infants may be evaluated and percentiles and larger databases would be added on these data shortly. In future studies infants will have only ultrasonographic measurements even if they do not have any additional intracranial pathology.

Another limitation is the timing of last ultrasonographic assessment which is at 34 weeks of gestational age. Our plan was to make a scan at 36 or 40 weeks of gestational age but this will be resulted in drop out of some infants. We did not change the discharge time of infants because of our study protocol and this would be an intervention and would not be ethical.

We do not know EABV is associated or related with neurologic outcome. We will follow up these patients and will be sharing neurodevelopmental results of our cohort as soon as possible.

Additionally there is no other clinican and/or radiologist doing all ultrasonographic measurements except ventricular index to assess an interobserver difference, this is also another limitation for this study.

## Conclusion

Anyway this method is safe, effective, easy to use and an applicable model for estimating preterm brain volume. As a conclusion, absolute brain volume could be calculated and estimated by 2D measurements obtained by transfontanel ultrasonography. All these measurements could be entered a database and/or computer base programme and may give more proper and reliable information about brain volume without any risk or workload in NICU even in low resource settings.

## Abbreviations

NHVM: Non hemorrhangic ventriculomegaly
c-PVL: Cystic periventricular leukomalacia
IVH: Intraventricular hemorrhage
NICU: Neonatal intensive care unit
ICH: Intracranial height
BPDia: Bi-parietal diamater
APD, FOD: Anteroposterior diamater or fronto-occipital distance
VI: Ventricular index
VH: Ventricular height
AHW: Anterior horn width
TOD: Thalamo-occipital distance
2D: Two dimensional
3D: Three dimensional
EABV: Estimated absolute brain volume
MRI: Magnetic resonance imaging
CrUS: Cranial ultrasonography

## Declarations

a. Ethics approval and consent to participate: All study approvals (Local instutional review board (Zekai Tahir Burak Hospital Specialty Educational Review Board (TUEK), Ethical board (Zekai Tahir Burak Hospital Clinical Ethics Committee, parental consents, this was in verbal form because cranial ultrasonography is a routine procedure during hospitalization in NICU) received before the initiation of this study.
b. Consent for publication. We have approval for publication of these data.
c. The datasets used and/or analysed during the current study are available from the corresponding author on reasonable request.
d. We have no any competing interests.
e. We did not receive any funding fort his study.
f. Authors’ contributions: GKS, FEC collected data and search literatüre, FEC made USG measurements, MB collected data and helped patient care, GKK, CT revised and approved the data.

## Notes

**Financial Disclosure:** None

